# OzTracs: Optical Osmolality Reporters Engineered from Mechanosensitive Ion Channels

**DOI:** 10.1101/2022.02.10.479916

**Authors:** Thomas J. Kleist, I Winnie Lin, Sophia Xu, Grigory Maksaev, Mayuri Sadoine, Elizabeth S. Haswell, Wolf B. Frommer, Michael M. Wudick

## Abstract

Interactions between physical forces and membrane proteins underpin many forms of environmental sensation and acclimation. Microbes survive sudden osmotic stresses with the help of mechanically gated ion channels and osmolyte transporters. Plant mechanosensitive ion channels have been shown to function in defense signaling. Here, we engineered genetically encoded osmolality sensors (OzTracs) by fusing green fluorescent protein spectral variants to the mechanosensitive ion channels MscL from *E. coli* or MSL10 from *A. thaliana*. When expressed in yeast cells, OzTrac sensors reported osmolality changes as a proportional change in emission ratio of the two fluorescent protein domains. Live-cell imaging revealed accumulation of fluorescent sensors in internal aggregates presumably derived from the endomembrane system. Thus, OzTrac sensors serve as osmolality-dependent reporters through an indirect mechanism, such as effects on molecular crowding or fluorophore solvation.

## Introduction

Organisms experience diverse mechanical forces arising from their internal or external environments. Internally, cells and cellular components generate forces during cell division, elongation, and movement. Externally, both unicellular and multicellular organisms face constantly changing environments that frequently present mechanical and osmotic stresses. Soil-dwelling organisms experience extreme cycles of osmolality changes due to, for example, soil drying in the sun or rehydration during rainfall ^1^. To be able to sense and respond to such changes, organisms evolved mechanosensors that monitor and respond to membrane tension changes. Mechanosensitive (MS) ion channels embedded in membranes are capable of detecting mechanical forces in membranes ^2,3^. MS ion channels have been broadly categorized as either tethered (*e*.*g*., to the cytoskeleton or cell wall / extracellular matrix) or intrinsically sensitive to membrane tension. Channels gated directly by membrane tension play a critical role when, for example, unicellular organisms such as bacteria must maintain membrane integrity when confronted with osmotic stresses. During hypoosmotic stress, bacterial MS channels open directly in response to increased membrane tension and mediate ion efflux causing a passive extrusion of water from the cytosol and resulting in a reduced cellular volume ^4,5^. These “osmotic safety valves” mediate solute efflux during hypoosmotic shock to protect cells from lysis caused by excess membrane tension ^6^. Bacterial MS channels, especially MscL and MscS (mechanosensitive channel of large and small conductance, respectively), have been characterized extensively. Structurally, *Ec*MscL (MscL from *E. coli*) and homologs form nonselective ion channels as pentamers with two transmembrane (TM) helices ^7,8^. *EcM*scS assembles as a homo-heptamer, with each subunit containing a short N-terminal periplasmic domain, three TM helices per subunit, and a C-terminal cytoplasmic region 9,10.

The MscS family is much larger than the MscL family and includes homologs from eukaryotes, unlike the MscL family. The MscS family is more variable in size and sequence across different species than the MscL family ^11^. Much of the diversity in the MscS family stems from variation in the number of TM helices per subunit, ranging from three to eleven. A family of ten MscS-like channels (MSL) in Arabidopsis (*Arabidopsis thaliana*) exhibits discernible structural homology to the pore and adjacent β-domain of *Ec*MscS but also contains diverse domains and topologies outside the pore-lining domain and adjacent regions ^12^. Some of the biological functions of eukaryotic MscS homologs have been characterized. For example, Msy1 and Msy2 from *Schizosaccharomyces pombe* localize to the endoplasmic reticulum (ER) membrane, where they influence cytosolic calcium (Ca^2+^) elevation in response to hypoosmotic shock ^11^. In Arabidopsis, MscS-Like 1 (*At*MSL1), whose structure has been recently solved by cryo-electron microscopy ^13^, localizes to the inner mitochondrial membrane where it is suggested to contribute to the dissipation of the mitochondrial membrane potential under abiotic stress ^14^. *At*MSL2 and *At*MSL3, as well as MSC1 from the alga *Chlamydomonas reinhardtii* ^15^, localize to the plastid envelope, where they may provide stretch-activated channel activity gated in response to osmotic imbalance between stroma and cytoplasm ^5,12,16^. The plasma membrane (PM)-localized *At*MSL8 is a pollen-specific membrane tension–gated ion channel essential for male fertility and required for pollen survival under hypoosmotic stress associated with rehydration ^17^. Arabidopsis MSL9 and MSL10 localize to the PM, and their gating properties have been studied in root protoplasts ^5^ and *Xenopus laevis* oocytes ^18^. Nonetheless, *msl9;msl10* double mutants and *msl4;msl5;msl6;msl9;msl10* quintuple mutant plants, which each lacked measurable stretch activated channel activity, responded normally to mechanical and osmotic stresses and showed no obvious phenotypic defects ^5^. Recently, *At*MSL10 was shown to function in cell swelling responses ^19^, transduction of oscillatory mechanical signals (Tran et al., 2021), and long-distance wound signaling ^20^. *At*MSL10 has been posited to act as a hydraulic sensor in plant vascular bundles, but the precise mechanism for *At*MSL10 activation during wound signaling is not fully understood.

Genetically encoded fluorescent biosensors have served as transformative tools for biologists in recent decades. Placement of sensory domains or full-length proteins between fluorescent protein domains spectrally compatible for Förster resonance energy transfer (FRET) has proven a facile and effective strategy for engineering genetically encoded biosensor prototypes ^21,22^. We chose a variety of mechanosensitive membrane proteins and used a sensor plasmid library from a prior study ^23^ to generate candidate sensors for expression in yeast (*Saccharomyces cerevisiae*). Unlike many soluble proteins, sensors constructed from integral membrane proteins are challenging to purify and work with *in vitro*. Thus, to screen for candidate membrane mechanosensors, live yeast cells expressing fluorescent chimeras were treated with solutions of varying osmotic potential and were monitored by fluorescence spectroscopy. Identified osmolality tracking or ‘OzTrac’ sensors based on *Ec*MscL or *At*MSL10 reported dose-dependent osmolyte treatments and were further characterized by trapped-cell microfluidics and confocal fluorescence microscopy, revealing sensor reversibility upon osmolyte removal. The *At*MSL10-derived OzTrac sensor retained mechanosensitive ion channel activity when expressed in frog (*Xenopus laevis*) oocytes, demonstrating functionality of the OzTrac chimera and its presence in the PM. In contrast, OzTracs aggregated internally when expressed in yeast, where they nonetheless function as sensors for osmotic potential. We hypothesize that the sensor mechanism is unrelated to membrane tension but rather caused by molecular crowding and/or changes in fluorophore solvation.

## Materials and methods

### Plasmid constructs

The full-length open reading frames (ORFs) of MSL10 (At5g12080), AHK1 (At2g17820) and OSCA1.1 (At4g04340) from *A. thaliana* and MscL (JW3252) from *E. coli* were cloned into the TOPO GATEWAY Entry vector. The yeast expression vectors were created by GATEWAY LR reactions between pTOPO plasmids and the pDRFLIP-GW yeast expression vector series ^23^, carrying fluorescent proteins of a FRET pair flanking the Gateway cassette (Table S1) following manufacturer’s instructions. For assays in *Xenopus laevis* oocytes, cDNAs were cloned into the oocyte expression vector pOO2-GW ^18^.

### Expression of sensors in protease-deficient yeast

The protease deficient yeast strain BJ5465 (*MATa ura3– 52 trp1 leu2Δ1 his3Δ200 pep4::HIS3 prb1Δ1*.*6R can1*) was obtained from the Yeast Genetic Stock Center (University of California, Berkeley, CA). Transformation of yeast cells was performed using a lithium acetate method ^24^. Transformants were selected on synthetic media containing yeast nitrogen base (YNB, Difco) supplemented with 2 % glucose and DropOut supplements lacking uracil (Clontech, Mountain View, CA). Single colonies were used to inoculate 5 mL liquid YNB media supplemented with 2 % glucose and DropOut supplements lacking uracil. Cells were grown with agitation (230 rpm) at 30 °C overnight until an OD_600nm_ of 0.2 - 0.3 was reached. Liquid cultures were sub-cultured by dilution to an OD_600nm_ of 0.1 in the same medium and grown at 30 °C with agitation until cultures reached OD_600nm_ ∼ 0.4. Cells were harvested by centrifugation for further analysis.

### Fluorimetry

Experiments were performed similarly to previous publications ^23,25,26^. Briefly, fresh yeast cultures (OD_600 nm_ ∼0.4) were washed three times in 50 mM 2-(*N*-morpholino)-ethanesulfonic acid (MES) pH 5.5, and resuspended in 50 mM MES pH 5.5. Fluorescence was measured with a fluorescence plate reader (M1000; TECAN, Austria) in bottom reading mode using 7.5 nm bandwidths for both excitation and emission ^27,28^. To quantify fluorescence responses of the sensors to different osmolytes, 100 µL aqueous solutions containing 50 mM MES pH 5.5 were added to 100 µL of cell suspension in 96-well flat-bottom plates (#655101; Greiner, Monroe, NC). Osmolyte titration curves were analyzed using the GraphPad Prism software. Data represent means and standard errors of the mean (SEM) of three technical replicates. Curves were fitted to a non-linear one-site specific binding with Hill slope and a constant: Y = B_max_ (X^h^)/(K_m_^h^ + X^h^) + a, with B_max_ being the maximum specific binding, K_m_ the Michaelis constant, h the Hill slope, which describes the cooperativity of the ligand binding, and a constant (a) added to the equation.

### Electrophysiology

*Xenopus laevis* oocytes were injected with complementary RNAs (cRNAs) encoding OzTrac-MSL10-34 that were amplified from linearized pOO2-GW-OzTrac-MSL10-34 plasmid. Excised inside-out patches were bathed symmetrically in 60 mM magnesium chloride (MgCl_2_), 10 mM 4-(2-hydroxyethyl)-1-piperazineethanesulfonic acid (HEPES), pH 7.4, and were subjected to pressure ramping from 0 to -140 millimeters of mercury (mmHg). The electrodes pulled from glass capillaries had a final resistance of 3-4 MOhm. The experiments were typically performed within 3-5 days after oocytes injection ^29^.

### Fluorescence Microscopy

Quantitative imaging was performed on a spinning disk confocal microscope. Yeast cells were trapped as a single cell layer in a microfluidic perfusion system (Y04C plate, Onyx, CellASIC, Hayward, CA, USA) and perfused with media containing 50 mM MES pH 5.5 with or without specified osmolytes. Microscopy data were acquired using an Olympus IXplore SpinSR confocal microscope equipped with UPLSAPO 100x 1.35 NA silicone immersion oil objective (Olympus), 50 µm spinning disk, and dual Photometrics Prime BSI sCMOS cameras. Acquisitions were performed using 2×2 pixel binning. A 75 mW 445 nm OBIS LX laser was used as an excitation source for DxDm and DxAm acquisitions, and a 40 mW coherent OBIS LX 514 nm laser was used to excite the AxAm channel. A 445/514/640 nm dichroic and 514 nm long pass beam splitter were used. For DxDm acquisition, a 482/35 nm emission filter was used, and a 534/23 nm emission filter was used for DxAm and AxAm acquisitions. A Nano-ZL300-OSSU fast piezo stage (Mad City Labs, Madison, Wisconsin) was used for z-stack acquisition, and an IX3 ZDC2 (Olympus) was used for z-drift compensation. Average z-stack projections were performed in ImageJ ^30^ prior to data analysis and figure preparation.

### Yeast growth assays

The *S. cerevisiae* strains used for hypoosmotic stress assays were BY4743 (*MATa/α his3Δ1*/*his3Δ1 leu2Δ0*/*leu2Δ0 LYS2*/*lys2Δ0 met15Δ0*/*MET15 ura3Δ0*/*ura3Δ0*) and the HomoDip knock out *fps1Δ* mutant in the same genetic background (YLL043W, clone ID: 31531, GE Dharmacon). Yeast cells were transformed with OzTrac-MscL36 or OzTrac-MSL10 using a lithium acetate-based method ^24^, and transformants were selected on synthetic medium lacking uracil (Clontech, Mountain View, CA) at 30°C. For hyperosmotic culture conditions, yeast cells were grown in liquid medium supplemented with 1 M sorbitol. For hypoosmotic shock treatments, yeast cells were transferred to liquid medium without sorbitol ^31^.

### Structural representations

Modeling of the structures of sensors was performed using UCSF Chimera software (UCSF, San Francisco, CA)^30^. Protein structures were obtained from RCSB Protein Data Base (PDB): CFP (2WSN), YFP (1YFP), *Mycobacterium tuberculosis* MscL (*Mt*MscL, 2OAR), and the three-dimensional structure of *At*MSL10 was generated in a previous study ^20^.

## Results and Discussion

### Construction and characterization of membrane mechanosensors

In organisms equipped with cell walls such as bacteria, fungi, and plants, the plasma membrane (PM) is typically under tension caused by turgor pressure.

Mechanical forces in the PM can therefore be affected by changing extracellular osmotic potential. Specifically, membrane tension can be reduced by treatment with hyperosmotic extracellular solutions or increased by treatment with hypoosmotic extracellular solutions. With the aim of constructing a membrane tension sensor, we fused FRET-compatible variants of cyan and yellow fluorescent proteins with integral membrane proteins reported to function as mechanosensitive, PM-localized proteins (Figure 1, Figure S1, Table S1, S2).

**Figure 1.**
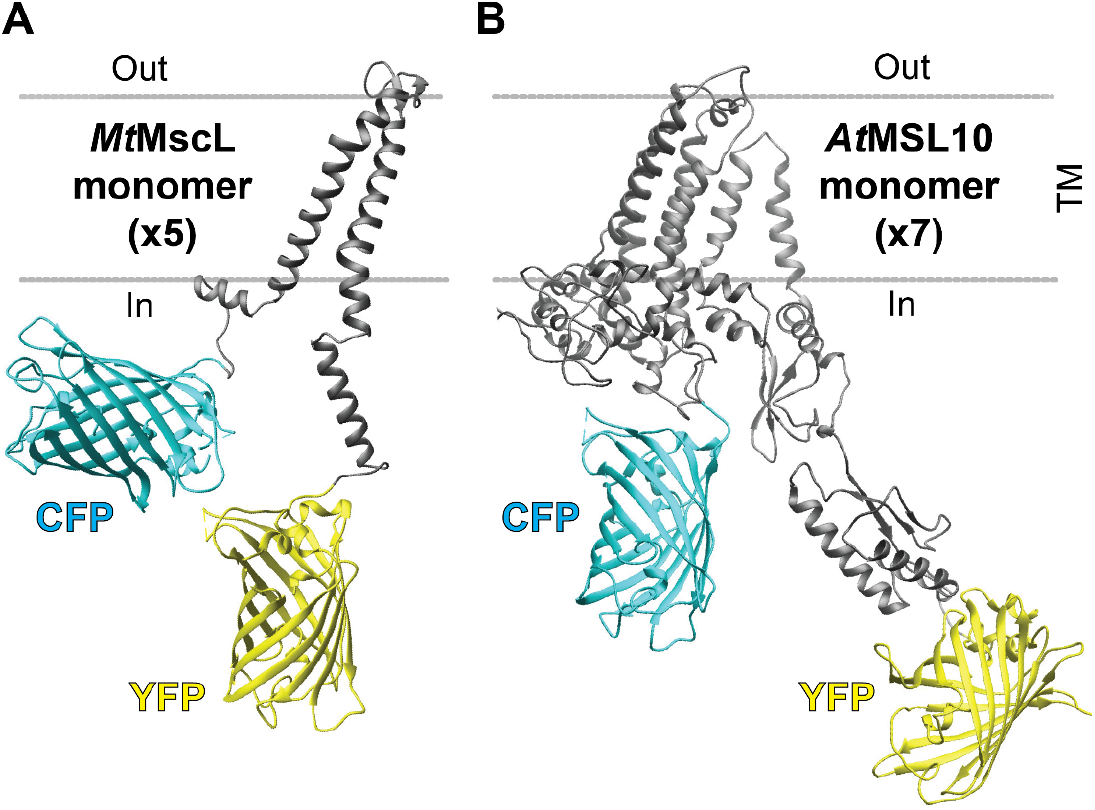
Monomeric structural representations of OzTrac sensors. Sensors were engineered by cloning the coding sequences of *Mt*MscL (A) and *At*MSL10 (B) between an N-terminal cyan fluorescent protein (CFP) and C-terminal yellow fluorescent protein (YFP). *Mt*MscL and *At*MSL10 are predicted to assemble as pentamers and heptamers, respectively. Visualization of the structure was performed using UCSF Chimera software and the three-dimensional structure of *At*MSL10 was generated in a previous study. TM: Trans-membrane domain.

Candidate sensory domains included the mechanosensitive ion channels *Ec*MscL and *At*MSL10. Chimeric sensors were expressed in *S. cerevisiae* cells, and fluorescence from live cells was monitored under hyperosmotic stress or isosmotic control conditions. OzTrac sensor responses were estimated as a change in the ratio of putative FRET acceptor emission (Am) to putative FRET donor emission (Dm) under FRET donor excitation (Dx): DxAm/DxDm. As the largest ratio changes were observed for *Ec*MscL- and *At*MSL10-based sensors, further characterization focused on those variants.

### Characterization of sensors constructed with *Ec*MscL

OzTrac candidates based on *Ec*MscL showed the largest DxAm/DxDm ratio changes with 3-45% greater DxAm/DxDm ratios when treated with 1 M sodium chloride (NaCl, 2 Osm/L) (Table S1). Direct effects on the acceptor fluorophore (i.e., acceptor emission under direct acceptor excitation (AxAm)) were not substantial (Figure 2A, Figure S2A, Table S1). Within the *Ec*MscL sub-group, the Aphrodite-Cerulean fluorophore combination (OzTrac-MscL-36) yielded the largest response with a +50 % ratio change (DxAm/DxDm) when cells were treated with 1 M NaCl, and was selected for further investigation in yeast.

**Figure 2.**
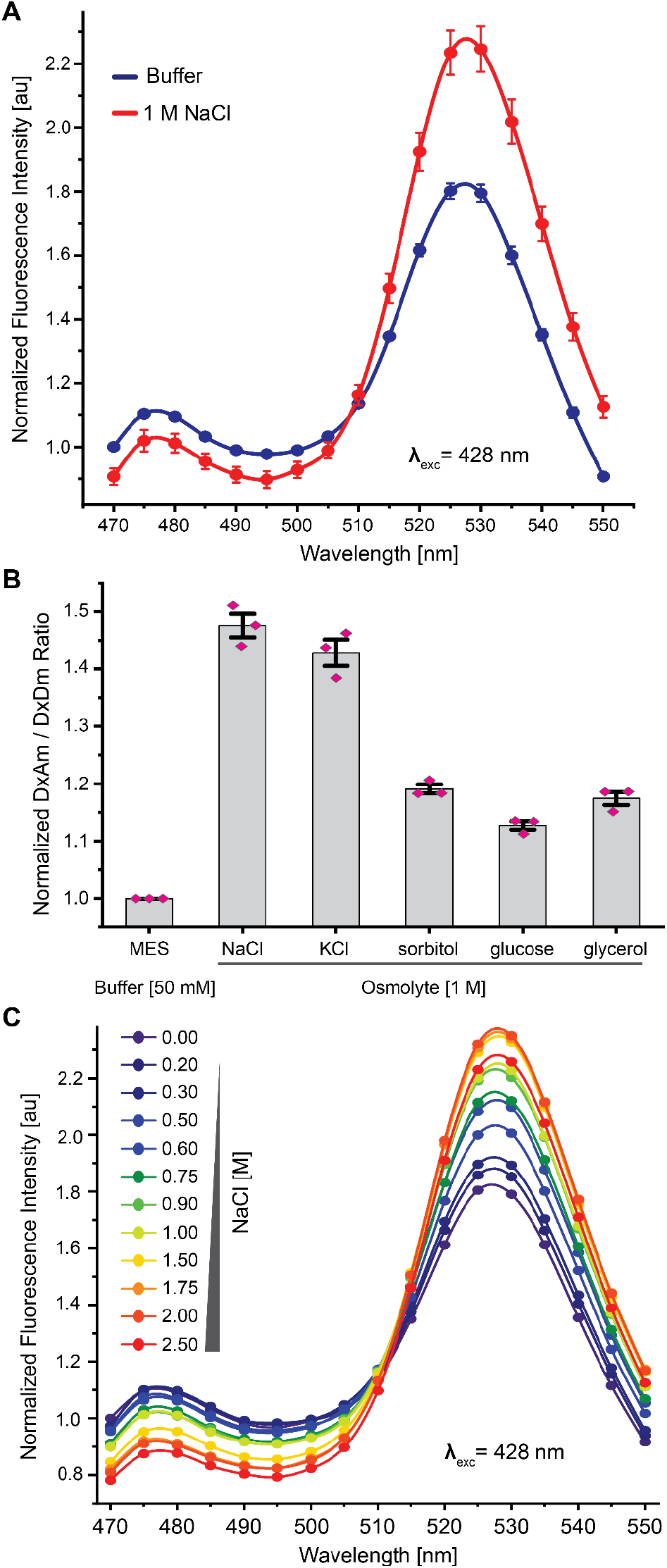
Characterization of the OzTrac-MscL-36 sensor. (A) Emission spectra of yeast cells expressing OzTrac-MscL-36 exposed to buffer with or without NaCl. X-axis: emission wavelength in nanometers (nm). Y-axis: fluorescence intensity normalized to 470 nm emission under control conditions. Excitation wavelengths (λ_exc_) are shown. (B) Various osmolytes, such as NaCl, KCl, sorbitol, glucose, or glycerol elicit ratiometric changes approximately proportional to solution osmolality. (C) Concentration-dependent effects of osmolyte (NaCl) treatment. Axes and inset same as in Figure 1A. All error bars: SEM.

To exclude that the observed ratio change could be caused by ion-specific effects on either the fluorophores or cellular host, other osmolytes were tested at 1 M concentration, including potassium chloride (KCl), sorbitol, glycerol, and glucose (Figure 2B). The addition of 1 M (2 Osm/L) KCl triggered a ratio change similar to that found for equimolar NaCl addition (∼45 %). Addition of 1 M (1 Osm/L) sorbitol, glucose or glycerol elicited lower ratio changes (+ ∼15-20 %). To estimate the dynamic response range of OzTrac-MscL-36, yeast cells expressing OzTrac-MscL-36were treated with NaCl concentrations ranging from 0-2.5 M and concentration-dependent spectral changes were observed for treatments with up to 1.75 M NaCl before reaching saturation (Figure 2C, Figure S2B), suggesting OzTrac-MscL-36 can report changes in extracellular osmotic potential over a wide range.

### Characterization of sensors constructed with *At*MSL10

Plants lack discernible MscL homologs but are equipped with homologs of MscS. Previous work had characterized Arabidopsis MSL10 by patch clamp in root protoplasts and oocytes, and had shown that it localized to the PM ^5,18^. In excised membrane patches, *At*MSL10 can be opened by application of positive or negative pipette pressure; channel closure occurs upon release of pressure. With the aim of creating a membrane tension sensor suitable for deployment in plants, we engineered fluorescent sensors based on *AtMSL10*, which was cloned into destination vectors containing 13 different fluorophore pairs ^23^ (Table S1), and named the candidates with the two largest response ratio changes (Aphrodite.t9-t7.mTFP.t9 and Aphrodite.t9-mTFP.t9) as OzTrac-MSL10-34 and OzTrac-MSL10-35, respectively.

Each sensor exhibited a ∼70% increase in ratio change when treated with 1 M NaCl (Figure 3A, Figure S2C, Table S1). Because of the similar responses and our interest in deployment in plants, further sensor characterization focused only on OzTrac-MSL10-34.

**Figure 3.**
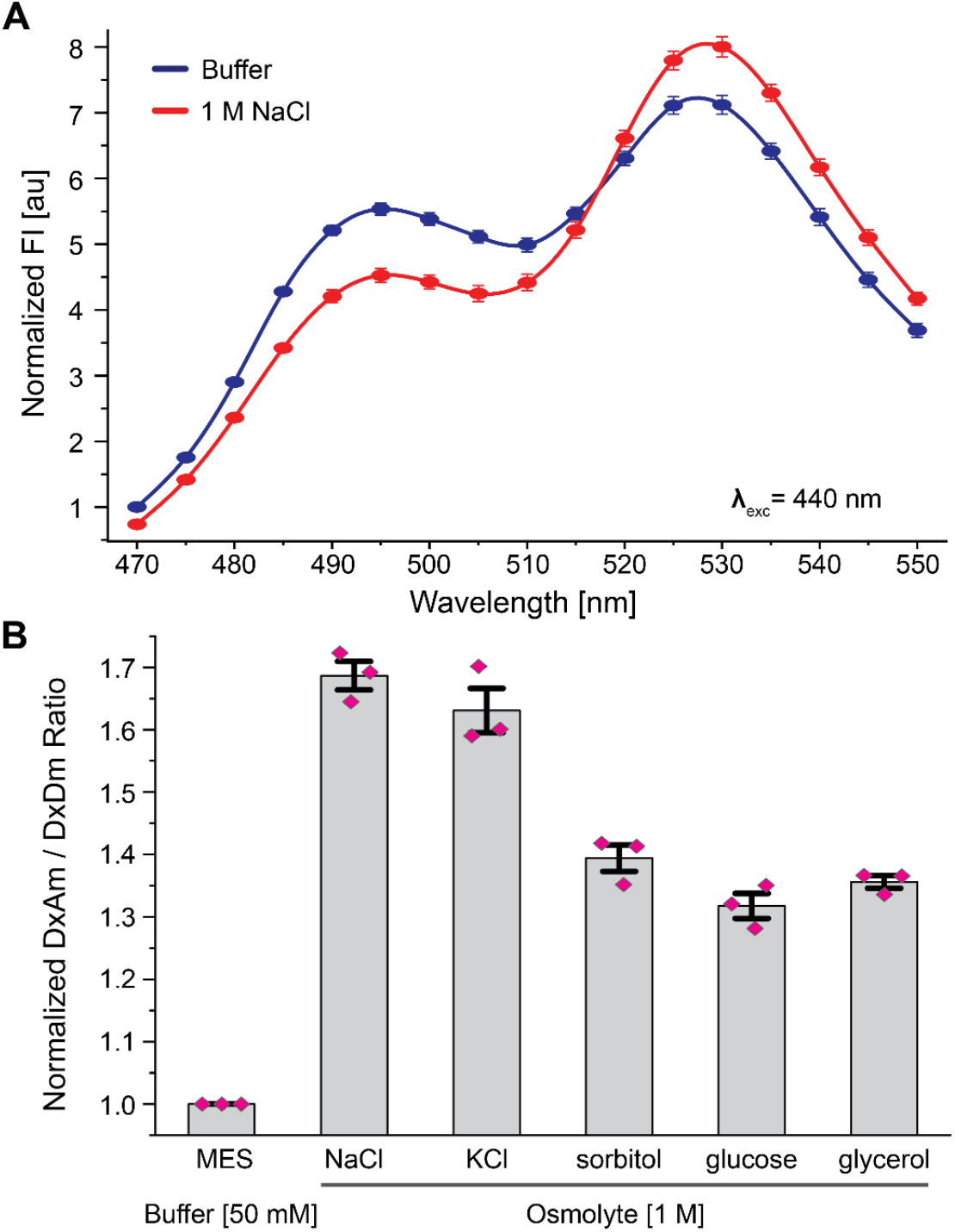
Characterization of the OzTrac-MSL10-34 sensor. (A) Emission spectra of yeast cells expressing OzTrac-MSL10-34 exposed to buffer with or without NaCl. (B) Response to osmolytes: NaCl, KCl, sorbitol, glucose, or glycerol (DxAm/DxDm) elicit ratiometric changes approximately proportional to solution osmolality. Axes and formatting as in Figure 2.

Similar to OzTrac-MscL-36, diverse osmolytes were able to elicit OzTrac-MSL10-34 DxAm/DxDm ratio changes (Figure 3B). Addition of 2 Osm/L KCl showed similar ratio DxAm/DxDm changes as NaCl (60-70% increase), whereas addition of 1 Osm/L sorbitol, glucose, and glycerol yielded smaller changes in ratio (30-40% increase) compared to NaCl and KCl, respectively.

Treatment of cells containing OzTrac-MSL10-34 with a concentration gradient of NaCl ranging from 0 - 1.88 M revealed concentration-dependent spectral differences (Figure 4A, Figure S2D). OzTrac-MSL10-34 effectively reported NaCl concentrations in the range of 200 - 800 mM with a dissociation constant (*K*_*d*_) of 573 ± 85 mM (Figure S3). Concentrations of NaCl > 0.8 M were associated with reduced fluorescence intensity from the acceptor fluorophore when exciting either the donor or the acceptor. Treatment with a gradient of glycerol concentrations ranging from 0- 2.5 M similarly revealed concentration-dependent spectral effects (Figure 4B, Figure S2E), with a working range from 0.32 - 1.8 M. *In planta, At*MSL10 functions as a stretch-activated ion channel that preferentially conducts anions ^18^. To test if stretch-activated channel activity is retained in the *At*MSL10-based OzTrac sensor, we characterized OzTrac-MSL10-34 by electrophysiology and yeast suppressor analysis.

**Figure 4.**
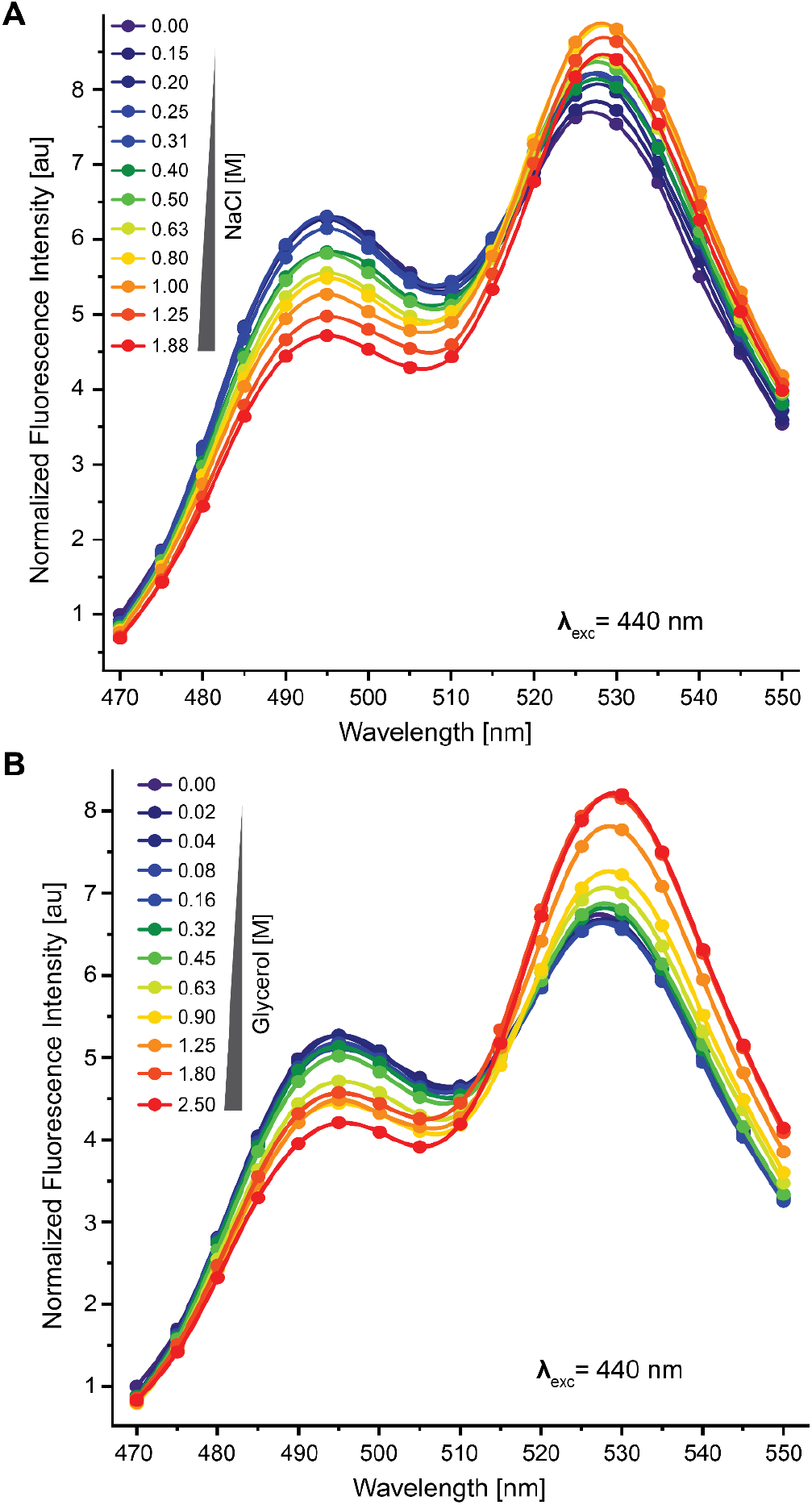
Osmolality-dependent ratio changes of OzTrac-MSL10-34 in yeast. (A) Response of the sensor to different NaCl concentrations. (B) Response to the sensor to different glycerol concentrations. Axes as in Figure 2.

### Electrophysiological analysis of OzTrac-MSL10-34

To test whether the OzTrac-MSL10-34 sensors is fully functional and forms stretch-activated ion channels, the chimera was expressed in oocytes and analyzed by patch clamping. Excised inside-out patches showed ion channel activity in response to pressure ramping (Figure 5). Unitary conductance appeared similar to previously reported values for *At*MSL10 ^18^, and gating pressure asymmetry (hysteresis), typical for wild-type *At*MSL10, was also observed. Together, the data indicate that OzTrac-MSL10-34 forms a functional ion channel, and that the fluorescent tags do not substantially alter *At*MSL10 channel properties or gating cycle.

**Figure 5.**
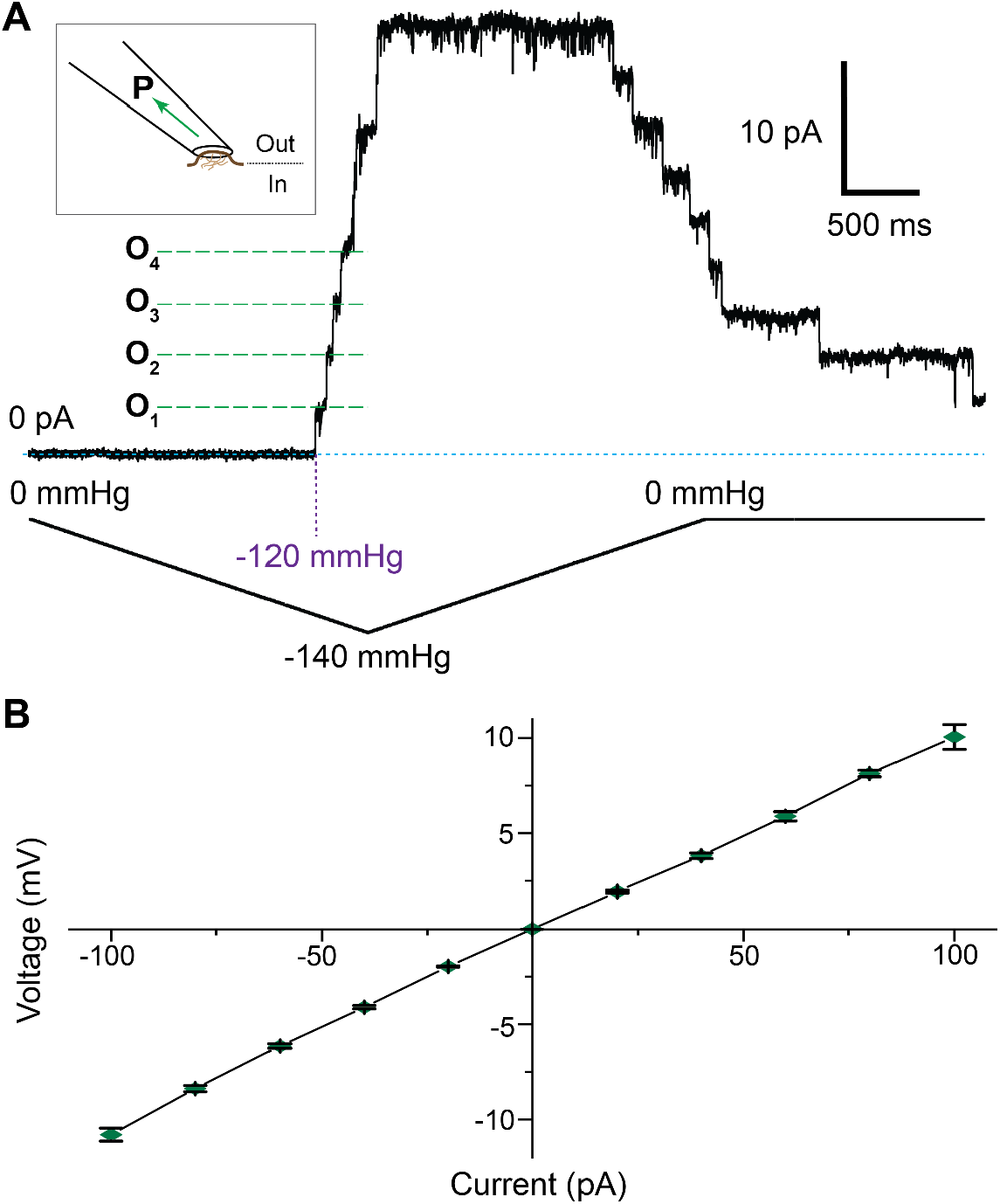
Ion channel activity of OzTrac-MSL10-34 expressed in *Xenopus laevis* oocytes. (A) Pressure ramp of an inside-out patch excised from oocytes. Schematic in upper left depicts excised patch of membrane; outer leaflet of the plasma membrane faces the interior of the pipette. Negative pressure (P) or suction was applied during patch clamp recordings. Example channel opening events (O_n_) are annotated and marked by green dotted lines. and pressure is given in millimetres of mercury (mmHg). Membrane potential = -40 mV. Note channel hysteresis, as previously described for *At*MSL10. (B) Current-voltage plot of currents observed in stretch-activated membrane patches. Unitary (single-channel) conductances were similar to published data for unmodified AtMSL10. Error bars: SEM.

#### Suppression of yeast fps1Δ mutant phenotype by expression of OzTracs

To test whether OzTrac-MscL-36 and/or OzTrac-MSL10-34 can function as ‘safety valves’ during hypoosmotic shock, as may be expected for functional mechanosensitive ion channels, the sensors were expressed in the yeast *fps1Δ* deletion mutant. The aquaglyceroporin Fps1 is required for survival in hypoosmotic shock conditions ^32^. Cells expressing the sensors were grown in liquid medium containing 1 M sorbitol, and serial dilutions of yeast cultures were inoculated onto solid medium with or without sorbitol. While the *fps1Δ* mutant grew on isosmotic medium containing 1 M sorbitol, little growth was observed on hypoosmotic media lacking sorbitol (Figure S4). Expression of either OzTrac-MscL-36 or OzTrac-MSL10-34 suppressed the *fps1Δ* phenotype by partially rescuing growth on media lacking sorbitol, consistent with the interpretation that OzTracs exhibit channel activity in yeast (Figure S4). It is unknown whether OzTrac-MSL10-34 or OzTrac-MscL derivatives transport glycerol, like FPS1, or if transport of other osmolytes may be responsible for the observed phenotypic suppression.

#### OzTrac-MSL10-34 likely reports molecular crowding or solvation status

To assess the subcellular localization and reversibility of sensor response, we turned to spinning disk confocal microscopy of trapped yeast cells.

Yeast expressing OzTrac-MSL10-34 were pressure-trapped in microfluidic devices and perfused with solutions of varying osmotic potential. Increased DxAm/DxDm ratios were elicited by treatment with 0.5 or 1.0 M sorbitol and could be reversed by osmolyte removal (Figure 6A). Strikingly, fluorescence was observed almost exclusively in internal structures that appeared as dense aggregates (Figure 6B).

**Figure 6.**
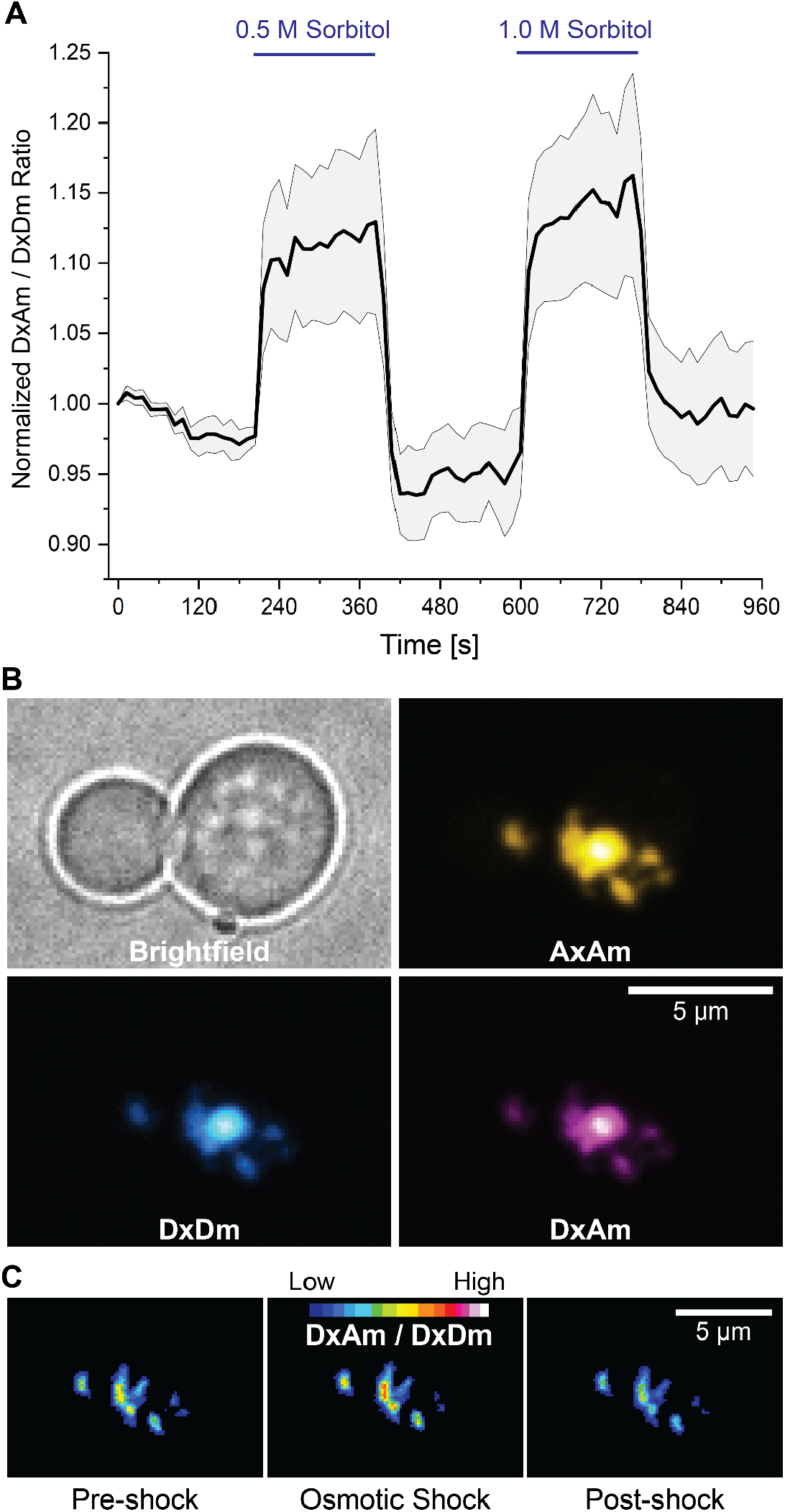
Quantitative fluorescence microscopy of yeast cells expressing OzTrac-MSL10-34. (A) Quantification of DxAm/DxDm ratio in trapped yeast cells exposed to square pulses of 0.5 or 1.0 M sorbitol (blue bars). Black line: mean (n = 5). Shaded region: SEM. (B) Example images of brightfield, AxAm, DxDm, and DxAm channels. Fluorescence images shown in “Hot” pseudocolor lookup tables (ImageJ). (C) Representative example of ratiometric (DxAm/DxDm) response from aggregates. 16-color lookup table applied. The same cell is shown in Figure 6B.

We surmise that sensor-containing aggregates arise from the endomembrane system and may be a consequence of overexpression of membrane proteins, as previously reported ^33^. Ratiometric imaging confirmed that changes in DxAm/DxDm ratio occurred in internal puncta (Figure 6C). Although we cannot exclude the possibility that a fraction of the sensor was present in the PM, our interpretation is that the OzTrac response is likely unassociated with events at the PM or membrane tension. Rather, we hypothesize that molecular crowding or differential hydration induced by hyperosmotic stress (as a consequence of changes in extracellular osmolality) may cause the observed response of intracellular OzTracs.

## Conclusions

In this study, we report construction of OzTracs, fluorescent sensors derived from mechanosensitive ion channels that report osmolality. OzTracs do not appear to target the cell membrane in yeast but rather aggregate internally. Localization to internal aggregates was unexpected given the suppression of the hypoosmotic shock-induced growth inhibition in the yeast *fps1Δ* strain. This apparent contradiction might be explained by the presence of a small population of sensors at the cell membrane below our fluorescence detection threshold. The mode of function for OzTracs is unclear, however we posit that the sensors undergo FRET changes caused by molecular crowding and/or changes in protein hydration status. Irrespective of the mechanism, OzTrac-MSL10-34 and OzTrac-MscL-36 constitute new tools for tracking external osmolality in yeast. Future work should include efforts to improve trafficking to the PM, for example by nested insertion of one or more fluorescent protein domains within the channel moiety. This approach has been successfully employed for development of sugar sensor-transporter chimeras that target the yeast PM ^34^. Given that we observed stretch-activated channel activity of OzTrac-MSL10-34 at the PM of *Xenopus* oocytes, the sensor may be suitable for use in animal and plant cell membranes.

## ASSOCIATED CONTENT

### Supporting information

Structural representations of multimeric *Mt*MscL or *At*MSL10; Acceptor fluorophore emission spectra under direct excitation; Hill slope fitting; Expression of pDRFLIP36 vector OzTrac-MscL-36 and OzTrac-MSL10-34 in the yeast plasma membrane aquaglyceroporin mutant (*fps1Δ*) cell line (BY4743); Table summarizing characterization of fluorophore pairs in pDRFLIP destination vector; Table of candidate proteins for construction of a fluorescent membrane tension sensor.

## Author Contributions

Conceptualization: WBF, Methodology: TJK, IWL, GM, ESH, WBF. Formal analysis: TJK, IWL, MS, GM. Investigation: TJK, IWL, SX, GM. Writing: TJK, IWL, GM, ESH, WBF, MMW. Visualization: TJK, IWL, MS. Supervision: TJK, IWL, ESH, WBF, MMW. Project Administration: WBF. Funding Acquisition: ESH, WBF.

## Notes

No competing financial interests have been declared.

## ACKNOWLEDGMENT

The authors thank Dr. David W. Ehrhardt for helpful discussions, and Cosima Sies for technical assistance.

## SYNOPSIS TOC

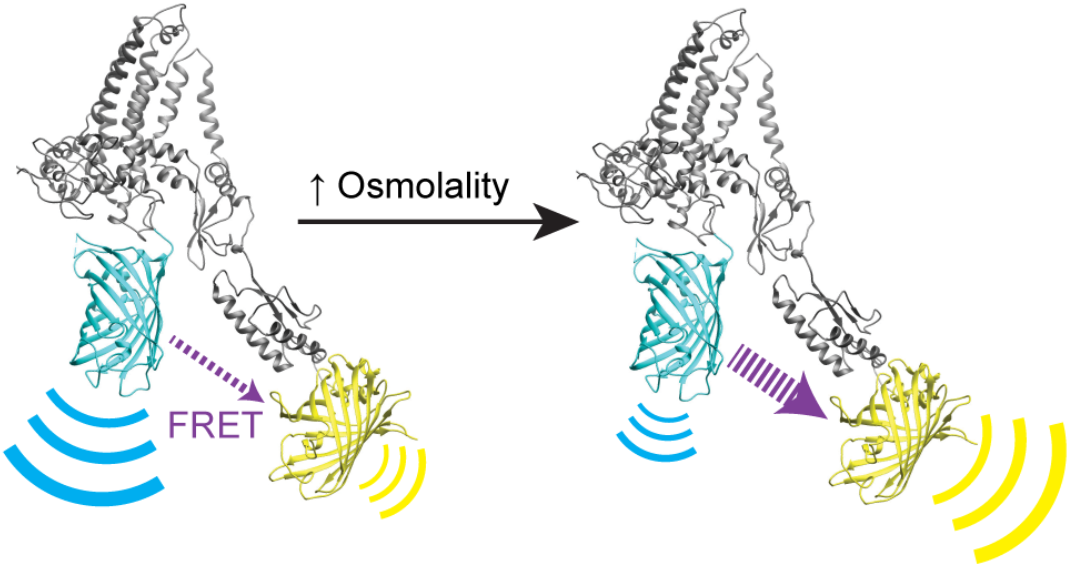

