## Supplemental figures and tables for "OzTracs: Optical Osmolality Reporters Engineered from Mechanosensitive Ion Channels"

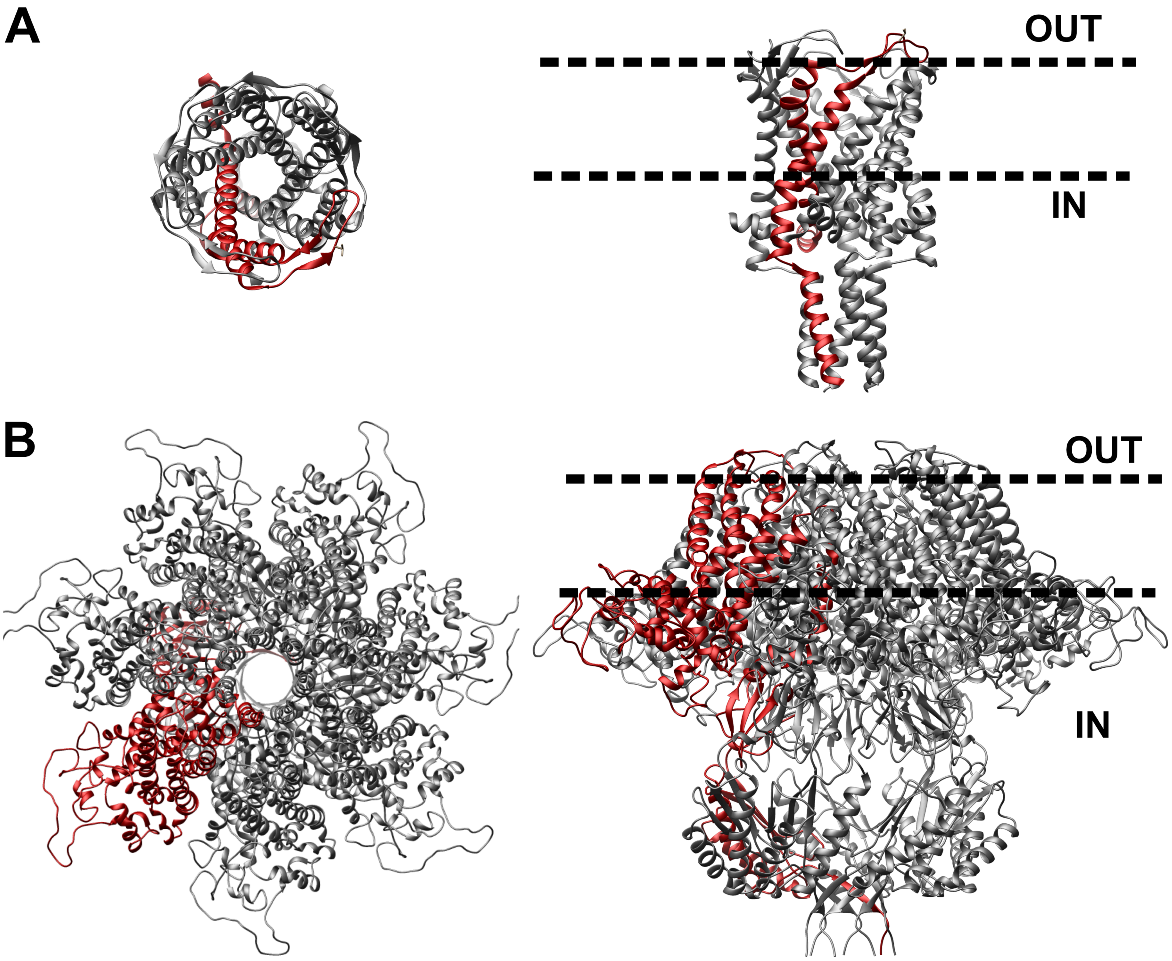


**Figure S1.** Structural representations of the multimeric assembly of (**A**) *Mt*MscL or (**B**) *At*MSL10 from top (left) and side (right) views. Single monomers are displayed in red. Visualization of the structure was performed using UCSF Chimera software, and the three-dimensional structure of *At*MSL10 was generated in a previous study [1].

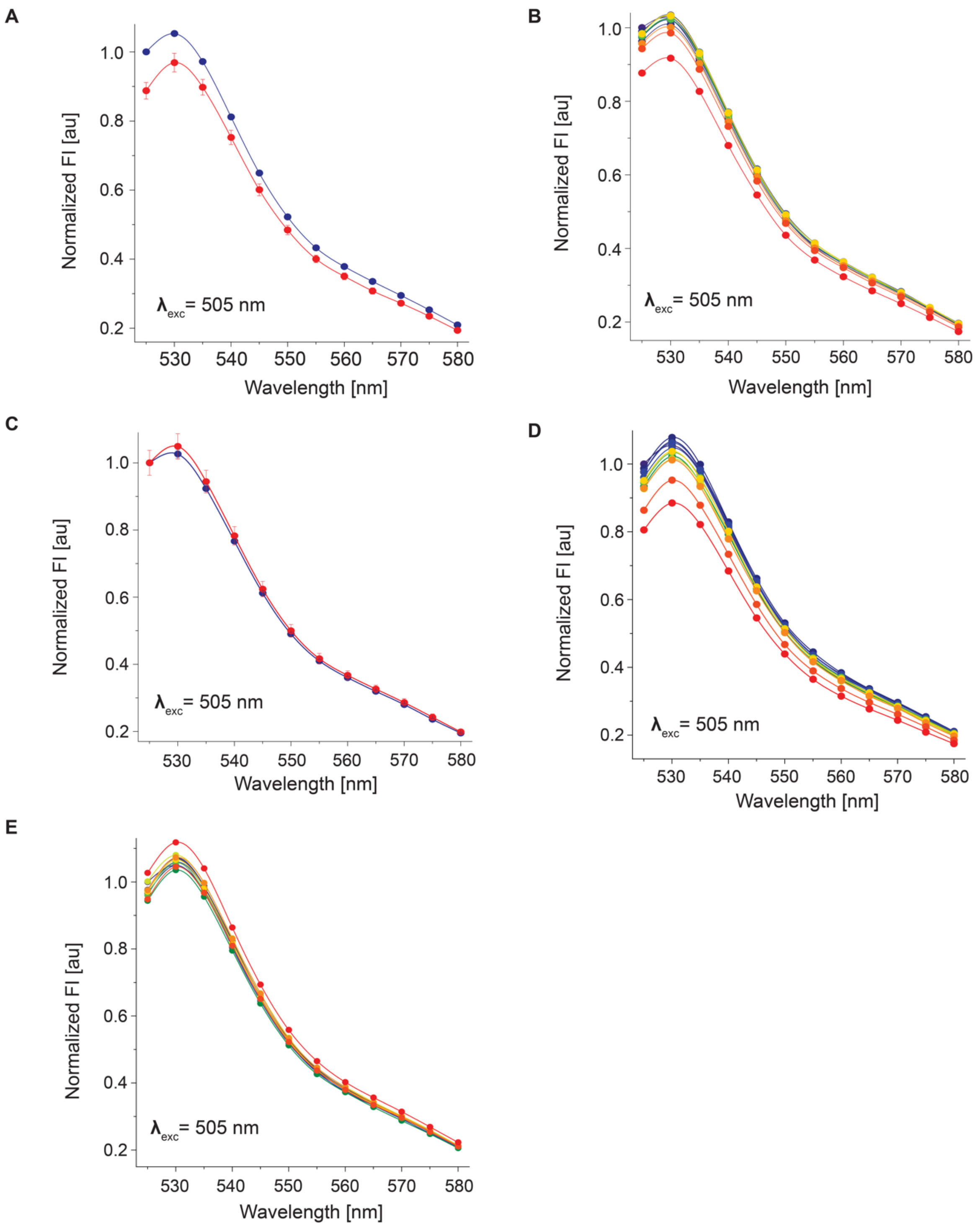


**Figure S2.** Acceptor fluorophore emission spectra under direct excitation (AxAm). **(A)** Emission spectra of yeast cells expressing OzTrac-MscL-36 exposed to buffer with (blue) or without NaCl (red) at 505 nm excitation. Control for Figure 2A. **(B)** Concentration-dependent effects of NaCl treatment on yeast expressing OzTrac-MscL-36 at 505 nm excitation. Control for Figure 2C. **(C)** Emission spectra of yeast cells expressing OzTrac-MSL10-34 exposed to buffer with (blue) or without NaCl (red) at 505 nm excitation. Control for Figure 3A. **(D)** Concentration-dependent effects of NaCl treatment on yeast expressing OzTrac-MSL10-34 at 505 nm excitation. Control for Figure 4A. **(E)** Concentration-dependent effects of glycerol treatment on yeast expressing OzTrac-MSL10-34 at 505 nm excitation. Control for Figure 4B.

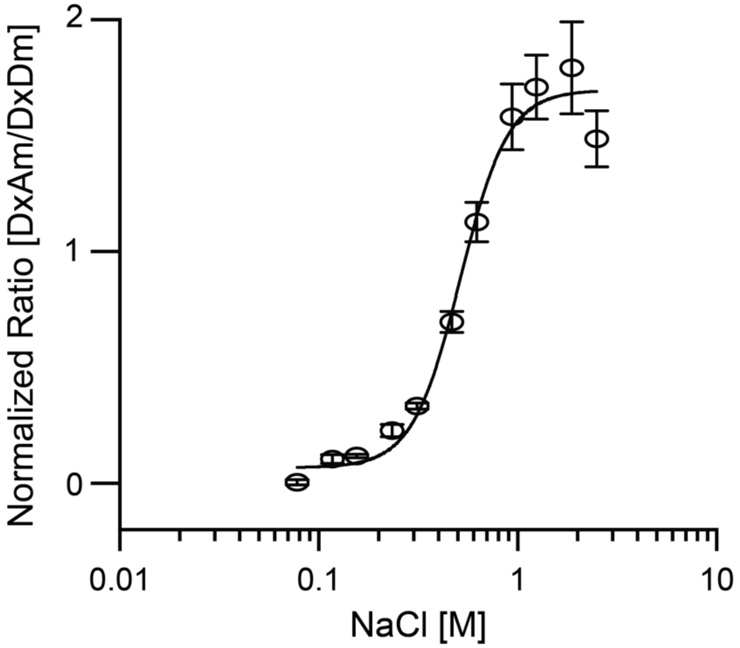


**Figure S3.** Ligand titration curve of OzTrac-MSL10-34 fitted to a non-linear one-site specific binding with a Hill slope, which describes the cooperativity of the ligand binding, and a dissociation constant ($K_d$) of 573 ± 85 mM.

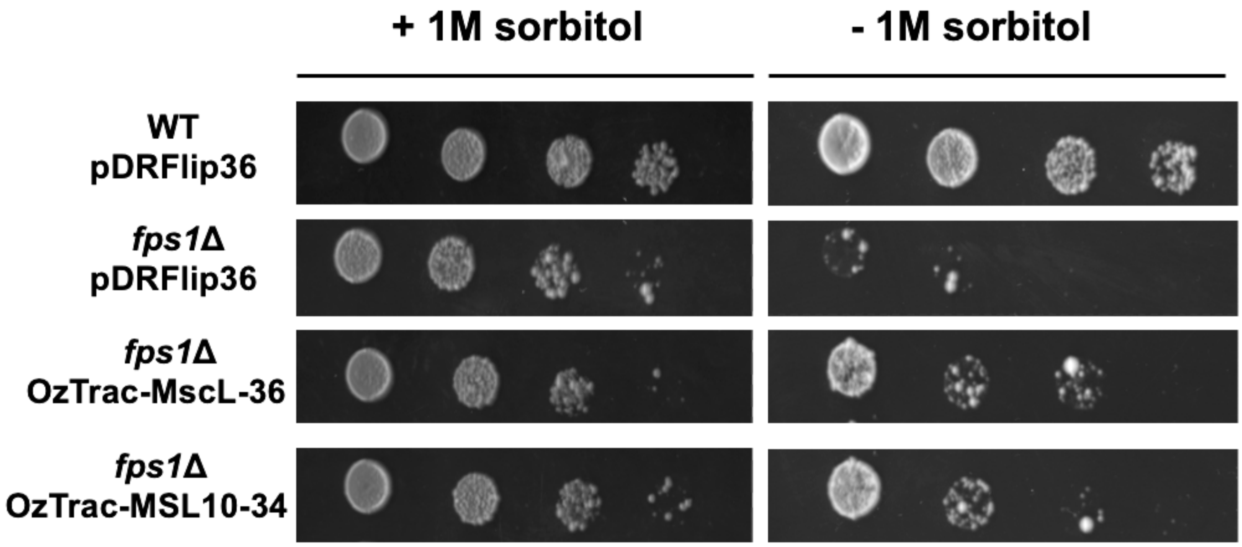


**Figure S4.** Partial suppression of the growth phenotype of the yeast plasma membrane aquaglyceroporin mutant *fps1Δ*. of expression of pDRFLIP36 (vector alone), OzTrac-MscL-36 or OzTrac-MSL10-34 in the yeast plasma membrane aquaglyceroporin mutant *fps1Δ* (BY4743). Cells were grown in liquid medium with 1 M sorbitol and then 1:10 serial dilution of yeast cultures were pipetted onto agar plates with (left panel) or without sorbitol (right panel) to expose cells to hypoosmotic conditions.

### SUPPLEMENTAL TABLES

**Table S1.** Fluorophore pairs in pDRFLIP destination vectors and observed DxAm/DxDm ratios when treated with 1 M NaCl. Higher values correspond to more intense red coloration.

| pDRFLIP # | FRET donor | FRET acceptor | DxAm/DxDm Response to 1 M NaCl | |
|---|---|---|---|---|
| | | | *Ec*MscL | *At*MSL10 |
| 30 | mCer | AFPt9 | 1.28 | 1.37 |
| 32 | t7CFPt9 | AFPt9 | 0.97 | 1.36 |
| 34 | t7TFPt9 | AFPt9 | ND | 1.65 |
| 35 | TFPt9 | AFPt9 | ND | 1.50 |
| 36 | Cer | AFPt9 | 1.45 | ND |
| 37 | Cer | Cit | 1.30 | ND |
| 38 | sCer | Cit | 0.90 | 1.00 |
| 39 | t7sCFPt9 | sAFPt9 | 1.03 | ND |
| 42 | mCer | Cit | ND | 1.52 |
| 43 | sCer | sAFPt9 | 1.20 | ND |
| 48 | mTrq2 | AFPt9 | 1.28 | ND |
| 49 | t7mTrq2t9 | AFPt9 | 1.20 | 1.40 |
| 50 | t7sTrq2t9 | sAFPt9 | ND | 1.20 |
| 51 | sTrq2 | sAFPt9 | ND | 1.05 |

*At* - *A. thaliana*, *Ec* - *E. coli*, ND - not determined.

**Table S2.** Candidate proteins for a FRET membrane tension sensor.

| **Candidate sensory protein (gene identifier)** | **Ref.** |
|---|---|
| *Ec*MscL (JW3252)<br>mechanosensitive channel of large conductance | 2 |
| *At*MSL10 (*At*5g12080)<br>mechanosensitive channel of small conductance-like 10 | 3 |
| *At*AHK1 (At2g17820)<br>protein histidine kinase 1 | 4 |
| *At*OSCA1.1 (At4g04340)<br>hyperosmolality-gated calcium-permeable channel 1.1 | 5 |

*At* - *A. thaliana*, *Ec* - *E. coli*, Ref – reference
